# Redox proteomics of stomatal immunity reveals role of a lipid transfer protein in plant defense

**DOI:** 10.1101/2020.05.21.108704

**Authors:** Kelly M Balmant, Sheldon R Lawrence, Benjamin V Duong, Fanzhao Zhu, Ning Zhu, Joshua Nicklay, Sixue Chen

## Abstract

Redox-based post-translational modifications (PTMs) involving protein cysteine residues as redox sensors are important to various physiological processes. However, little is known about redox-sensitive proteins in guard cells and their functions in stomatal immunity. In this study, we applied an integrative protein labeling method cysTMTRAQ and identified guard cell proteins that were altered by thiol redox PTMs in response to a bacterial flagellin peptide flg22. In total, eight, seven and 20 potential redox-responsive proteins were identified in guard cells treated with flg22 for 15, 30 and 60 min, respectively. The proteins fall into several functional groups including photosynthesis, lipid binding, oxidation-reduction, and defense. Among the proteins, a lipid transfer protein (LTP)-II was confirmed to be redox-responsive and involved in plant resistance to *Pseudomonas syringe pv. tomato* DC3000. This study not only creates an inventory of potential redox-sensitive proteins in flg22 signal transduction in guard cells, but also highlights the relevance of the lipid transfer protein in plant defense against the bacterial pathogens.

**Sentence summary:** Thiol-redox proteomics identified potential redox sensors important in stomatal immunity, and a lipid transfer protein was characterized to function as a redox sensor in plant immune response.

## INTRODUCTION

Pathogens cause serious damage to crops each year and threaten global food security. On average, yield losses due to pests and diseases range from 20% to 40% in major crops worldwide. In certain countries, pathogens can result in severe losses. For example, diseases caused up to 41% loss in rice in Asia, and 96% loss in potato in France (Cerda et al., 2017). Unable to penetrate the plants epidermis directly, bacterial pathogens rely on wounds and natural openings of plants to gain access and establish disease in plants. Stomatal pores dominate the aerial parts of plants and act as a first line of defense against bacterial pathogens. These tiny structures are composed of pairs of specialized epidermal cells called guard cells, which are responsible for regulating gas exchange and water loss through changing the aperture of stomatal pores (Balmant et al., 2016; Papanatsiou et al., 2017). For many years, stomata were thought to be a passive entrance for foliar bacterial pathogens. Now they are known to play an active role in limiting bacterial invasion as part of the plant innate immune system. Guard cells have the ability to perceive live bacteria as well as Pathogen Associated Molecular Patterns (PAMPs), and close the stomatal pores within one hour of exposure to the bacteria or PAMPs (e.g., the N-terminal 22 amino acid peptide from flagellin, flg22) (Melotto et al., 2006).

Pathogen perception initiates a signal transduction cascade where reactive oxygen species (ROS) and reactive nitrogen species (RNS) play important roles in activating downstream responses leading to stomatal closure (Vidhyasekaran, 2013; Medeiros et al., 2020; Wang et al., 2020). The production of ROS and RNS is a common event during stomatal closure (Pham and Desikan, 2009; Wang et al., 2020), and is one of the earliest events during plant response to bacterial pathogen. In fact, loss of cysteine thiol (S)-nitrosylation activity in *Arabidopsis* was shown to disable plant defense response conferred by resistance (R) genes. For example, dysregulation of S-nitrosylation blocked both salicylic acid (SA) signaling and SA synthesis (Feechan et al., 2005). In order to achieve rapid immune responses, plant cells employ such PTMs of key proteins in signaling pathways (Balmant et al., 2016). NONEXPRESSER OF PR GENES 1 (NPR1) is one of the few examples of redox-regulated proteins in plant defense. S-nitrosylation of Cys156 promotes NPR1 oligomerization. In the presence of pathogen, cellular redox changes mediated by SA lead to reduction of the NPR1 oligomer to monomers, which are transported into the nucleus (Mou et al., 2003; Waszczak et al., 2015). In the nucleus, SA-mediated redox changes cause reduction of TGA transcriptional factors so that they form an active complex with NPR1 to turn on pathogenesis related (PR) genes; NPR1 is subsequently phosphorylated, leading to ubiquitination and degradation (Spoel et al., 2009; Waszczak et al., 2015). In addition, it was shown that constitutively high S-nitrosothiol (SNO) levels promote a decrease in cell death through S-nitrosylation of Respiratory Burst Oxidase Homolog D (RBOH D), leading to reduction in its activity and oxidative stress (Yun et al., 2011). Furthermore, nitric oxide (NO) was shown to promote S-nitrosylation of SA-binding protein 3 (SABP3) in *Arabidopsis* during the NO burst, suppressing its ability to bind SA and decreasing its chloroplast carbonic anhydrase activity (Wang et al., 2009).

As described above for NPR1, RBOH D and SABP3, cysteines residues of certain proteins are sensitive to ROS and RNS (Hoshi and Heinemann, 2001). The high pKa values of cysteines make these residues sensitive to small redox perturbation by forming reactive ionized thiolate groups (Spoel and Loake, 2011). When exposed to oxidative stress, the thiol groups are capable of undergoing reversible inter- and intra-molecular disulfide bond formation, nitrosylation, sulfinic acid modification, glutathionylation, and irreversible sulfonic acid modification (Balmant et al., 2016; Lawrence et al., 2020). Although in the past few years, the knowledge of guard cells hormone signaling networks has improved (Zhao et al., 2010; Zhu et al., 2010, 2014; Balmant et al., 2016; David et al., 2019), the understanding of the guard cell innate immune response against bacterial invasion is very limited. Consequently, much remains unknown about the molecular components and regulations involved in the stomatal immune response. Furthermore, there is a growing interest to understand how PTMs regulate various aspects of guard cell innate immunity.

One crucial aspect in redox proteomics is the importance of addressing the issue of protein turnover during experimentation. Overlooking this important issue may lead to misleading results. In order to tackle this problem, we employed a double-labeling strategy called cysTMTRAQ (Parker et al., 2015), where the isobaric tags cysTMT and iTRAQ are employed in the same experiment for simultaneous determination of quantifiable cysteine redox changes and protein level changes. Using this powerful tool, we were able to create an inventory of previously unknown potential redox proteins and highlight some protein regulatory mechanisms in stomatal guard cell innate immunity. Among these proteins, we identified a lipid transfer protein (LTP)-II undergoing oxidation in response to flg22 during stomatal closure. LTPs are small, basic proteins present in higher plants. They are known to be involved in key cellular processes such as stabilization of membranes, cell wall organization, and signal transduction. LTPs are also known to play important roles in plant response to biotic and abiotic stresses, as well as in plant growth and development (Liu et al., 2015). Here, we report redox proteomics discovery of the LTP-II, its functional characterization, and present a potential mechanism by which LTP-II is involved in plant defense response.

## RESULTS

### Flg22 Induction of Stomatal Closure and ROS Production

To test whether flg22 induces stomatal closure in *B. napus* as in *Arabidopsis* and tomato (Melotto et al., 2006), epidermal peels were treated with different concentrations of flg22 and stomatal closure was examined. As shown in Fig. S1, 1 μM and 3 μM flg22 caused significant stomatal closure within 1 hour. As flg22 concentration increased to 10 μM, the effect of the treatment became more obvious. To ensure a significant effect of flg22 treatment within the two-hour period, a concentration of 10 μM flg22 was used in further experiments. This concentration has been applied in previous studies (Melotto et al., 2006; Wang et al., 2012; Lee et al., 2015). Enzymatically digested epidermal peels without the pavement and mesophyll cells are enriched for guard cells. The ability of enriched stomatal guard cells to respond to flg22 was also tested. Although the enriched stomata did not close as much as was observed in the epidermal peels, they were still highly responsive to flg22. As shown in Fig.1A, stomatal closure was significant after 30 min flg22 treatment. The use of enriched stomatal guard cells allows correlation of stomatal movement (as an immune response) in real-time with physiological and molecular changes in the guard cells.

ROS are known to play a central role in flg22-induced stomatal closure (Navarro et al., 2004; Pham and Desikan, 2009; Bigeard et al., 2015). Here we showed that 10 μM flg22 induced ROS production in *B. napus* enriched stomatal guard cells (Fig. 1B). The enriched stomatal samples showed an increase in H_2_O_2_ in response to flg22, with the highest peak at 15 minutes after flg22 treatment (Fig. 1B). These results demonstrate that *B. napus* enriched stomatal guard cells are responsive to flg22, and suggested that ROS and/or the guard cell redox state change may play important roles in the signal transduction leading to stomatal closure.

**Figure 1.**
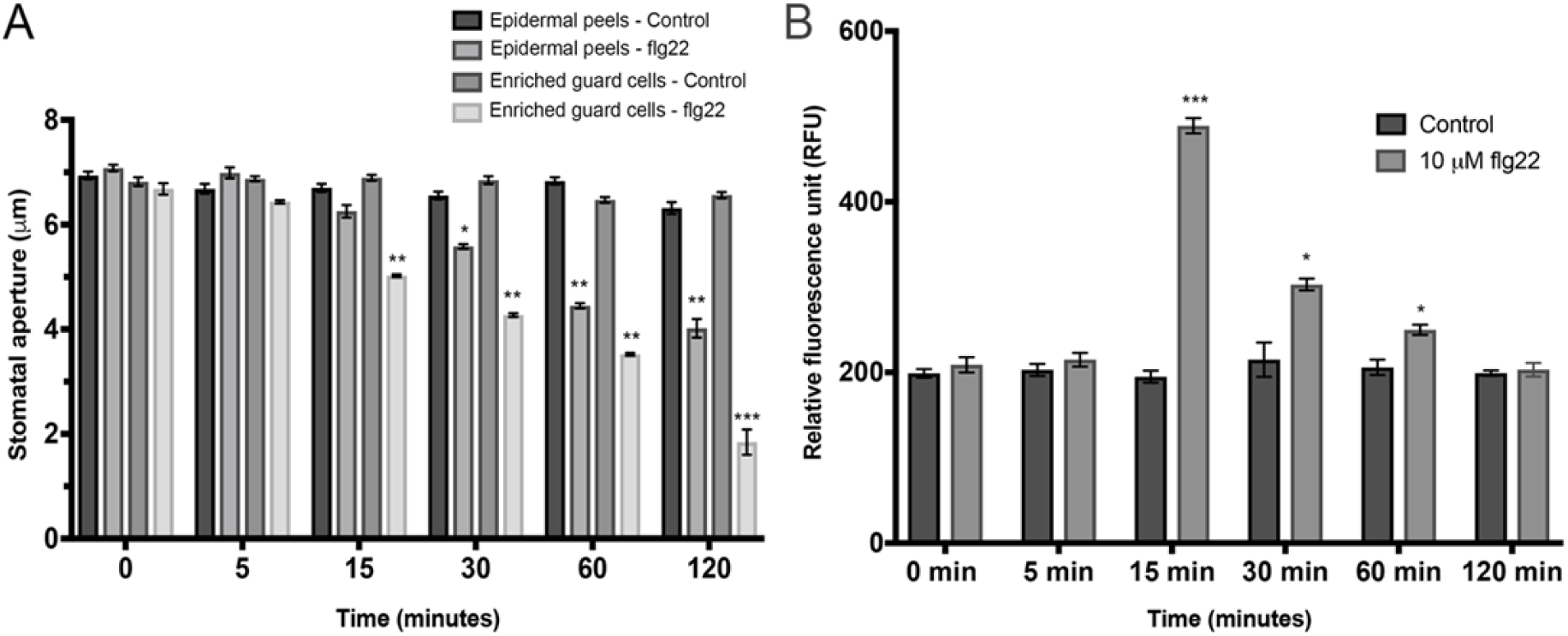
Stomatal closure and ROS generation in response to flg22. (**A**) Stomatal closure in response to 10 μM flg22 added at time 0 min in epidermal peels and enriched guard cells samples. Data were obtained from 0 stomata from three independent experiments and presented as means ± SE. (**B**) Enriched guard cells were treated with either water (Control) or 10 μM flg22 and subjected to ROS measurement assay. The data are shown as means ± SE of three independent experiments. The asterisks indicate significantly different mean value of bacterial growth between WT and *ltp-II* plants at p<0.05.

### Identification of Flg22-Responsive Proteins and Redox Proteins in Guard Cells

To investigate redox-sensitive proteins potentially involved in the flg22 triggered stomatal closure, we used a double labeling method cysTMTRAQ (Parker et al., 2015). Importantly, a reverse labeling procedure was performed, in which iodoacetamide was included in the protein extraction step to block free thiol groups. This reverse-labeling strategy maintains the initial redox state of the proteins and prevents artificial oxidation during sample preparation. Thus, increase of cysTMT signals from specific peptides derived from treated samples compared to control samples indicates the presence of oxidation responsive cysteine thiols. The use of this double-labeling strategy enabled identification of flg22-regulated proteins and redox-sensitive cysteine residues in the guard cells. In total, 399, 488 and 434 unique cysteine-containing peptides corresponding to 211, 221 and 209 proteins were confidently identified (FDR 0.05) among three biological replicates of the samples treated with 10 μM flg22 for 15, 30 and 60 min, respectively. To determine flg22-responsive redox proteins, we compared the relative levels of different cysteine-containing peptides (indicated by peak areas of the different cysTMT tags) in control and flg22-treated samples. A threshold of fold change greater than 1.2 or less than 0.8, and a q-value smaller than 0.05 were set as stringent criteria for significant differences between the control and treatment. A total of five, four, and 11 proteins underwent oxidation upon 15 min, 30 min and 60 min flg22 treatment, respectively (Table 1). Additionally, a total of three, three and nine proteins underwent reduction after flg22 treatment for 15 min, 30 min and 60 min, respectively (Table 1).

**Table 1.**
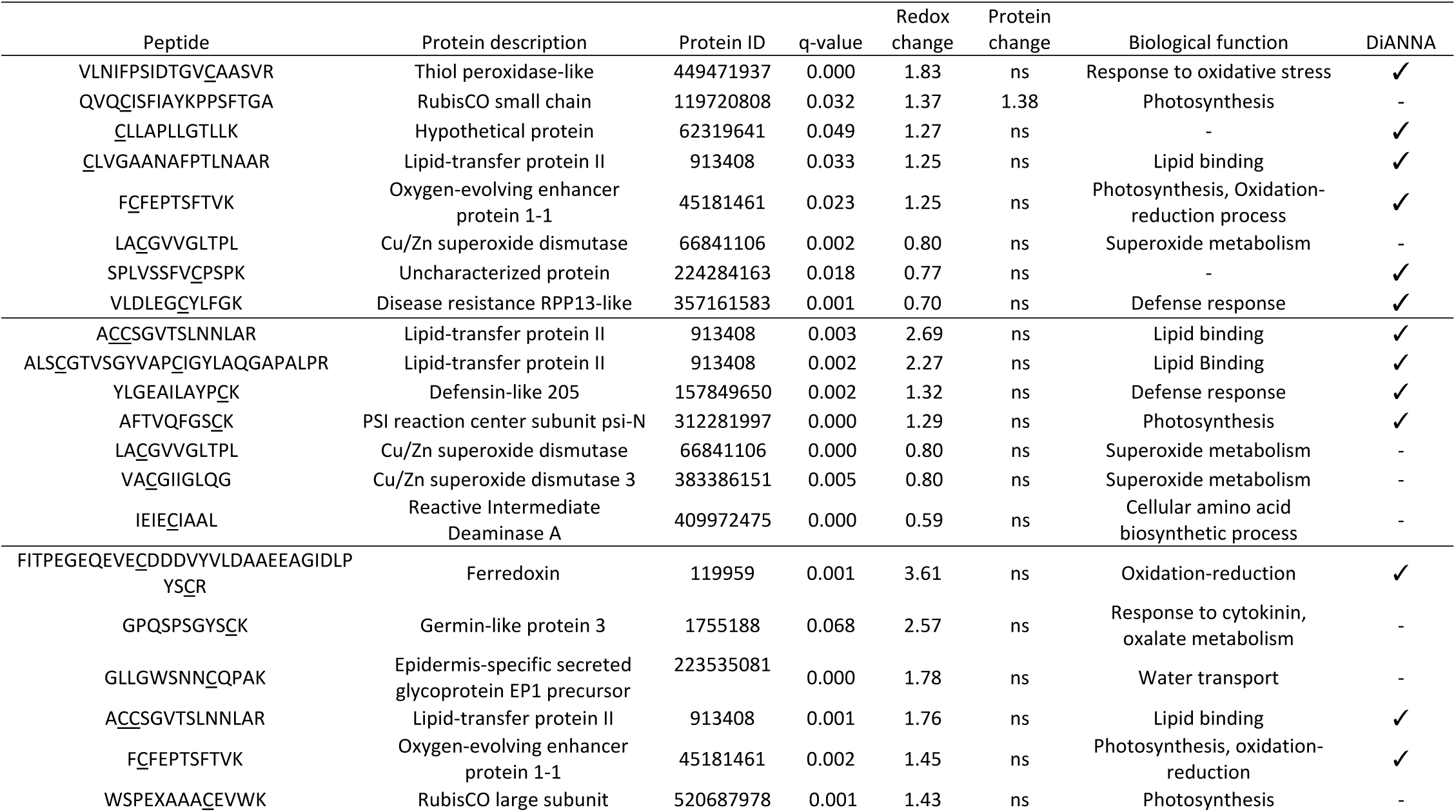
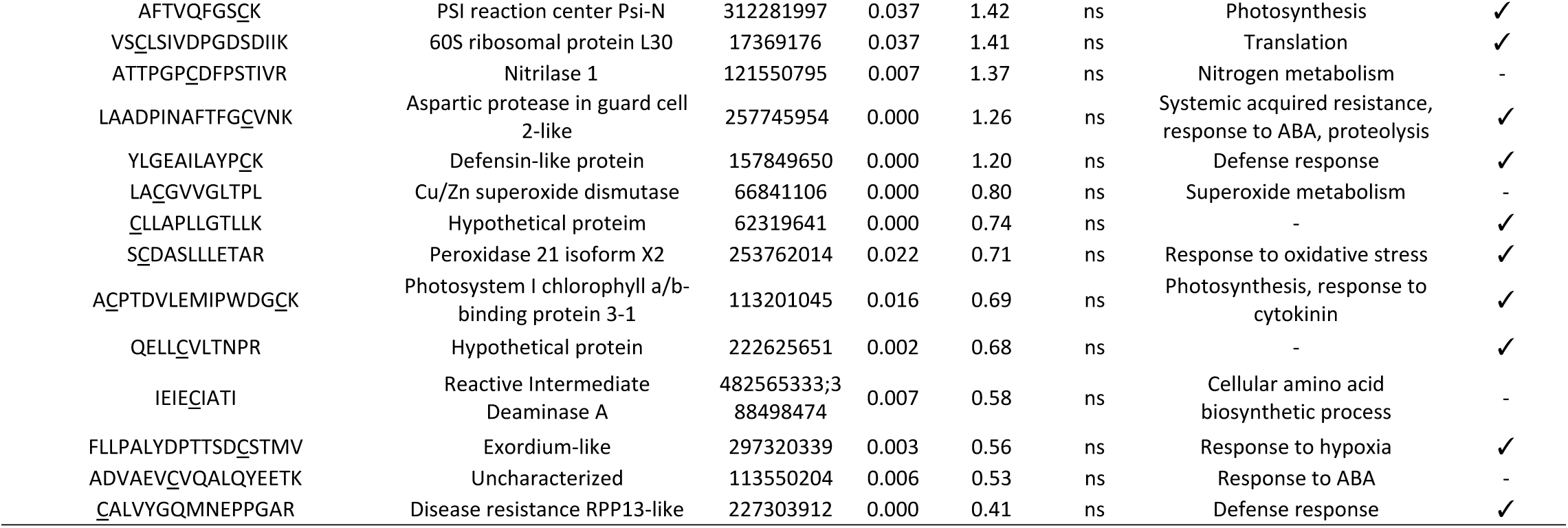
Redox-responsive proteins identified in *B. napus* guard cells after 15 min (top), 30 min (middle) and 60 min (bottom) treatment with flg22. The tick under DiANNA indicates the predicted disulfide bond formation. The q-value represents the p-value adjusted for multiple correction in the differential analysis.

As previously reported, identification of redox-regulated proteins may be complicated by possible protein level changes (Alvarez et al., 2009; Parker et al., 2015). Here we used information of total protein relative quantification (indicated by peak areas of the different iTRAQ tags) to overcome this issue. In total, 2734, 3082 and 3220 unique peptides corresponding to 1402, 1519 and 1586 proteins were confidently identified (FDR 0.05) among the three biological replicates treated with flg22 for 15 min, 30 min and 60 min, respectively. A total of nine, 15 and 35 proteins were observed with increased expression levels in the samples treated with flg22 for 15 min, 30 min and 60 min, respectively. In addition, a total of 16, seven and 13 proteins showed substantial decreases in abundance in the above samples, respectively (Table S1).

### Characteristics of Guard Cell Redox Proteomic Changes in Response to Flg22

A total of eight potential redox-regulated proteins were identified in guard cell at 15 minutes after exposure to flg22 (Table 1). Five proteins (hypothetical protein, lipid transfer protein II, oxygen-evolving enhancer protein 1, RuBisCO small subunit, and thiol peroxidase-like) were found to be in the oxidized state. When the total protein-level change was taken into account, one protein (RuBisCO small subunit) out of the eight-potential redox-regulated proteins also showed significant protein amount change in the same direction (Table 1). Therefore, the observed redox-fold change of RuBisCO small subunit may be due to the protein level change rather than the thiol redox response. Using DiANNA software (http://clavius.bc.edu/~clotelab/DiANNA/), seven of the eight redox-sensitive proteins were predicted to form intra-molecular disulfide bonds (Table 1). The GO biological processes of the potential redox-regulated proteins include lipid binding, photosynthesis, oxidation-reduction process, superoxide metabolic process, defense response, and response to oxidative stress.

At 30 minutes after exposure to flg22, seven cysteine-containing peptides from six different proteins were identified as being redox-sensitive (Table 1). Three of the potential redox-regulated proteins (lipid-transfer protein II, photosystem I (PSI) reaction center subunit psi-N, and defensin like protein 205) were found to be in oxidized states, while the other three proteins (reactive intermediate deaminase A, Cu/Zn superoxide dismutase (SOD) 2, and Cu/Zn SOD 3) were in reduced states compared to the mock control. It is important to note that lipid-transfer protein II and Cu/Zn SOD 2 were also found to be potentially redox-regulated at 15 minutes after flg22 exposure. None of these proteins showed significant protein abundance changes based on iTRAQ quantification. Their biological processes include lipid binding, cellular amino acid biosynthetic process, response to toxic substance, superoxide metabolic process, and defense response (Table 1).

At the late stage of stomatal closure, 20 cysteine-containing peptides from 20 different proteins were found be potentially redox-regulated (Table 1). Twelve (lipid transfer protein II, PSI reaction center subunit psi-N, nitrilase 1, oxygen-evolving enhancer protein 1, ferredoxin, epidermis-specific secreted glycoprotein EP1 precursor, germin-like protein, aspartic protease in guard cell 2-like, Cu/Zn SOD, 60S ribosomal protein L30, ribulose-1,5-bisphosphate carboxylase/oxygenase large subunit, and defensin-like protein 205) were in oxidized states, whereas the other eight (chlorophyll a/b-binding protein, uncharacterized protein, disease resistance protein RPP13-like, two hypothetical proteins, protein exordium-like, reactive intermediate deaminase A, and peroxidase 21) were in reduced states compared to the control samples (Table 1). It is worth mentioning that lipid-transfer protein II and Cu/Zn SOD 2 were identified as redox-regulated proteins in all the three time points. In addition, three proteins (PSI reaction center subunit psi-N, reactive intermediate deaminase A, and defensin-like protein 205) that were shown undergoing oxidation at 30 minutes after flg22 exposure were also shown to be potentially redox-regulated at the late stage of stomatal closure. Furthermore, oxygen-evolving enhancer protein 1 was found to undergo oxidation, and two other proteins (hypothetical protein and disease resistance protein RPP13-like) undergo reduction at 15 minutes after flg22 exposure. They were also shown to be potentially redox-regulated at the late stage of stomatal closure. None of the proteins exhibited significant protein amount changes (Table S1). A total of 13 potential redox-regulated proteins (out of 20) are predicted to form intra-molecular disulfide bonds. Biological processes of the 20 potential redox-regulated proteins include photosynthesis, response to cytokinin, response to ABA, defense response, oxidation-reduction process, response to hypoxia, cellular amino acid biosynthetic process, response to toxic substance, systemic acquired resistance, proteolysis, superoxide metabolic process, response to oxidative stress, and translation (Table 1).

### *ltp-II* Plants Show Enhanced Disease Susceptibility to *Pst* DC3000

The protein LTP-II was found to undergo oxidation at both early and late stages of stomatal closure in response to flg22 and therefore was chosen for further analysis (Table 1). The structure of plant LTPs contain eight cysteine residues located at conserved regions among different species, and the cysteine residues are known to form four disulfide bonds (Fig. S2A and S2C), creating an internal hydrophobic cavity, which contains the lipid-binding site (Yeats and Rose, 2008). LTPs are encoded by a multigene family in a diversity of plant species (Fig. S2B) and are usually divided into two classes, LTPa and LTPb, based on their molecular weight (Finkina et al., 2016). In *Arabidopsis*, LTP-II belongs to the LTPa class, which contains 15 members (Fig. S2D). They share amino acid sequence similarity ranging from 20% to 70%.

*A. thaliana* and *B. napus* are members of the *Brassicacea* and are phylogenetically close relatives that share many sequence similarities (Parkin et al., 2005). For example, the coding region of *AtLTP-II Arabidopsis* gene (GeneID: *AT2G38530*) and its homologue in *B. napus* (*BnLTP-II*, GeneID: *106450956*) are 79% similar at the protein level (Fig. S3). This close phylogenic relationship allows the use of *Arabidopsis* (a reference plant with enormous genetics resources) as a powerful tool to characterize the functions of the LTP-II identified in this study. Since some LTPs have been shown to be involved in plant resistance to biotic stress (see previous section), we reasoned that plants impaired in *LTP-II* expression may compromise their disease resistance response compared to wild type (WT) plants. To test this hypothesis, we dip-inoculated *Arabidopsis* WT and homozygous *ltp-II* mutant plants with the virulent bacterial pathogen *Pseudomonas syringae* pv *tomato* DC3000 (*Pst DC3000*) and examined the bacterial growth in the inoculated plants. As shown in Fig. 2A, the growth of *Pst* DC3000 in *ltp-II* plants was significantly higher than in WT plants, indicating that the LTP-II plays a positive role in plant resistance to biotic stress.

**Figure. 2.**
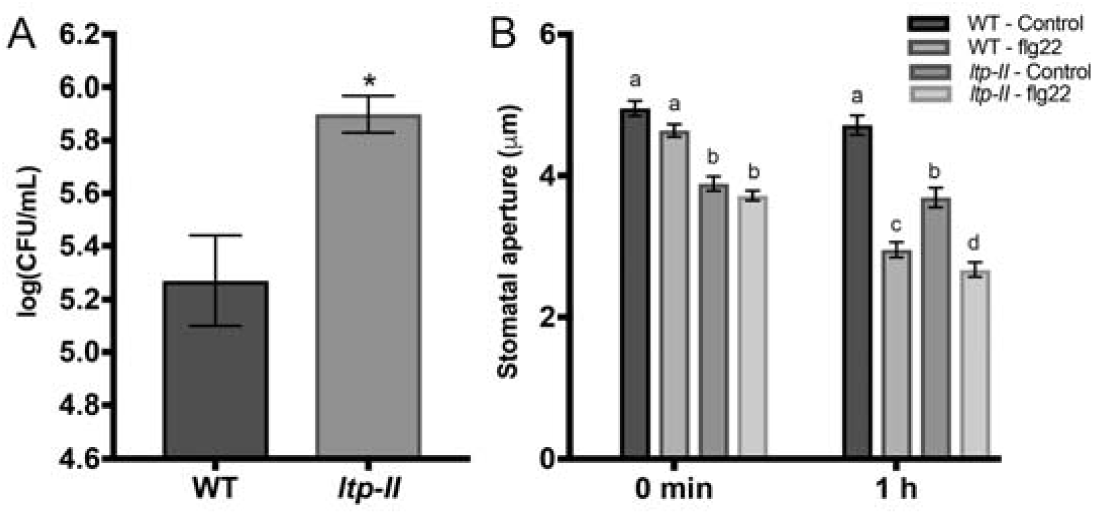
Responses of Wild-type (WT) and the *ltp-II* mutant to *P. syringae* and flg22. (**A**) Bacterial growth. WT and *ltp-II* mutant were inoculated with *Pst* DC3000. Samples were taken at 3-day post-inoculation (dpi) to determine the bacterial titers. (**B**) Stomatal aperture in response to 10 μM flg22. At least 180 stomata from three biological replicates were analyzed. The data are shown as means ± SE of three independent experiments.

### *ltp-II* Plants Show Smaller Stomatal Aperture than WT

The observation that *ltp-II* plants showed enhanced disease susceptibility to *Pst* DC3000 prompted us to hypothesize that the *ltp-II* plants had impaired stomatal closure, allowing more bacteria to enter and colonize the plants. To determine the *ltp-II* stomatal movement in response to biotic stress, epidermal peels were incubated in the opening buffer without (mock) and with 10 μM of flg22 for 1 hour under light. The stomatal apertures of WT and *ltp-II* plants were measured. To our surprise, the *ltp-II* plants had significantly smaller stomatal aperture than WT plants with or without flg22 treatment, and the stomatal movement was not impaired (Fig. 2B). These results suggest that stomatal aperture is not the main factor in the enhanced susceptibility of *ltp-II* plants.

### LTP-II Functions as a ROS Scavenger in Guard Cells

It is well established that ROS are second messengers in stomatal closure (Navarro et al., 2004; Pham and Desikan, 2009; Bigeard et al., 2015), and also known to play a major role in hypersensitive response of plant defense against pathogens (Baxter et al., 2014; Frederickson Matika and Loake, 2014). To examine the ROS levels in the *ltp-II* plants, we used a sensitive fluorophore dichlorofluorescein (H_2_DCF) to measure the changes of H_2_O_2_ levels in guard cells. In this assay, the nonpolar diacetate ester H_2_DCF-DA permeates into the guard cells, and is hydrolyzed in to polar and nonfluorescent H_2_DCF, which is trapped and subsequently oxidized by H_2_O_2_ to yield the fluorescent DCF (Zhang et al., 2001). As shown in Fig. 3A, exogenous application of flg22 enhanced the DCF fluorescence intensity in guard cells. Moreover, when compared to WT plants, the *ltp-II* plants showed higher DCF fluorescent intensity, indicating that these mutant plants have more ROS accumulation than WT (Fig. 3A). Interestingly, the *ltp-II* plants also showed higher ROS levels than WT even without the flg22 treatment (Fig. 3A). In addition to measuring the ROS levels, we quantified the contents of free thiols in the *ltp-II* and compared with WT plants. As expected, the contents of free thiols were significantly lower in WT plants after the flg22 treatment. Intriguingly, the mock control *ltp-II* plants showed a lower free thiol content than WT plants, and flg22 treated *ltp-II* plants had even lower free thiols (Fig. 3B). Altogether, these results support the hypothesis that *ltp-II* cells are under oxidative stress and LTP-II may play a role in ROS scavenging.

**Figure. 3.**
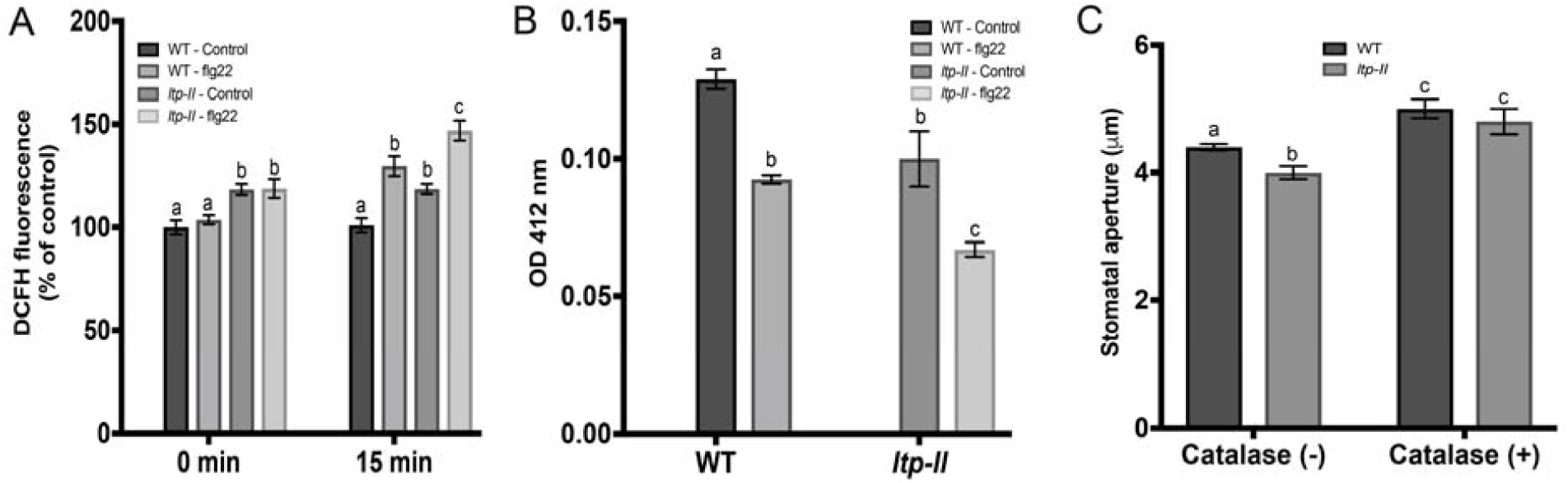
ROS accumulation and redox changes in WT and the *ltp-II* mutant in response to flg22. (**A**) flg22 caused increases in stomatal H2DCFDA fluorescence. Mean fluorescence level was significantly higher in *ltp-II* than WT. Fluorescence is relative to WT control samples at 0 min. (**B**) flg22 treatment perturbs cellular free thiol content, which was monitored using DTNB. The *ltp-II* plants showed more oxidation than WT plants in response to flg22. (**C**) Stomatal aperture in the presence of ROS scavenger catalase. In the presence of catalase, the *ltp-II* and WT plants exhibited the same size of stomatal aperture. At least 180 stomata from three biological replicates were analyzed. The data are shown as means ± SE of three independent experiments.

To test the hypothesis that LTPII is playing a role in ROS scavenging, we treated the epidermal peels of both WT and the *ltp-II* plants with 200 U/mL catalase (an enzyme known to convert H_2_O_2_ to H_2_O and O_2_) and analyzed stomatal aperture. After catalase treatment, there was no significant difference in stomatal aperture between the WT and *ltp-II* plants (Fig. 3C). These results indicate that the *ltp-II* is deficient in ROS scavenging and LTP-II functions as a ROS scavenger. The results also suggest that the smaller stomatal aperture observed in the *ltp-II* plants (Fig. 2B) may be due to its high ROS levels.

### *BnLTP-II* complements the *ltp-II* mutant of *Arabidopsis*

To further confirm the role of LTP-II in ROS scavenging/antioxidant activity, and *BnLTP-II* as the functional ortholog of *AtLTP-II*, we expressed *BnLTP-II* in the *Arabidopsis ltp-II* mutant (Fig. S5). First, to investigate the role of LTP-II in plant resistance to biotic stress, we examined bacterial growth in the leaves of WT Col-0, *ltp-II mutant*, and two independent lines of *LTP-II* overexpressing plants *LTP-OE8* and *LTP-OE12* (Fig. 4A; Fig. S5). Three days after dip-inoculation (3 dai) with *Pst DC3000*, bacterial growth was significantly increased in the *ltp-II* line compared to WT and the overexpressing lines, confirming that LTP-II plays a positive role in plants response to biotic stress (Fig. 4A). Clearly, complementation with *BnLTP-II* was adequate to rescue the mutant disease phenotype. Second, we examined the role of LTP-II in stomata movement in response to biotic stress. Epidermal peels of WT, *ltp-II mutant, LTP-OE8* and *LTP-OE12* plants were incubated with 10 μM of flg22 and stomatal apertures were measured after one hour. Surprisingly, at as early as 15 min flg22 treatment, *ltp-II mutant* plants showed significantly smaller stomatal aperture compared to WT and *LTP-OE8* and *LTP-OE12*. Complementation with *BnLTP-II* was able to rescue the mutant phenotype (Fig. 4B). These results confirm that stomatal aperture is not the casual factor in the enhanced susceptibility of *ltp-II* plants. Importantly, they demonstrate the critical function of the LTP-II in stomatal immunity.

**Fig. 4.**
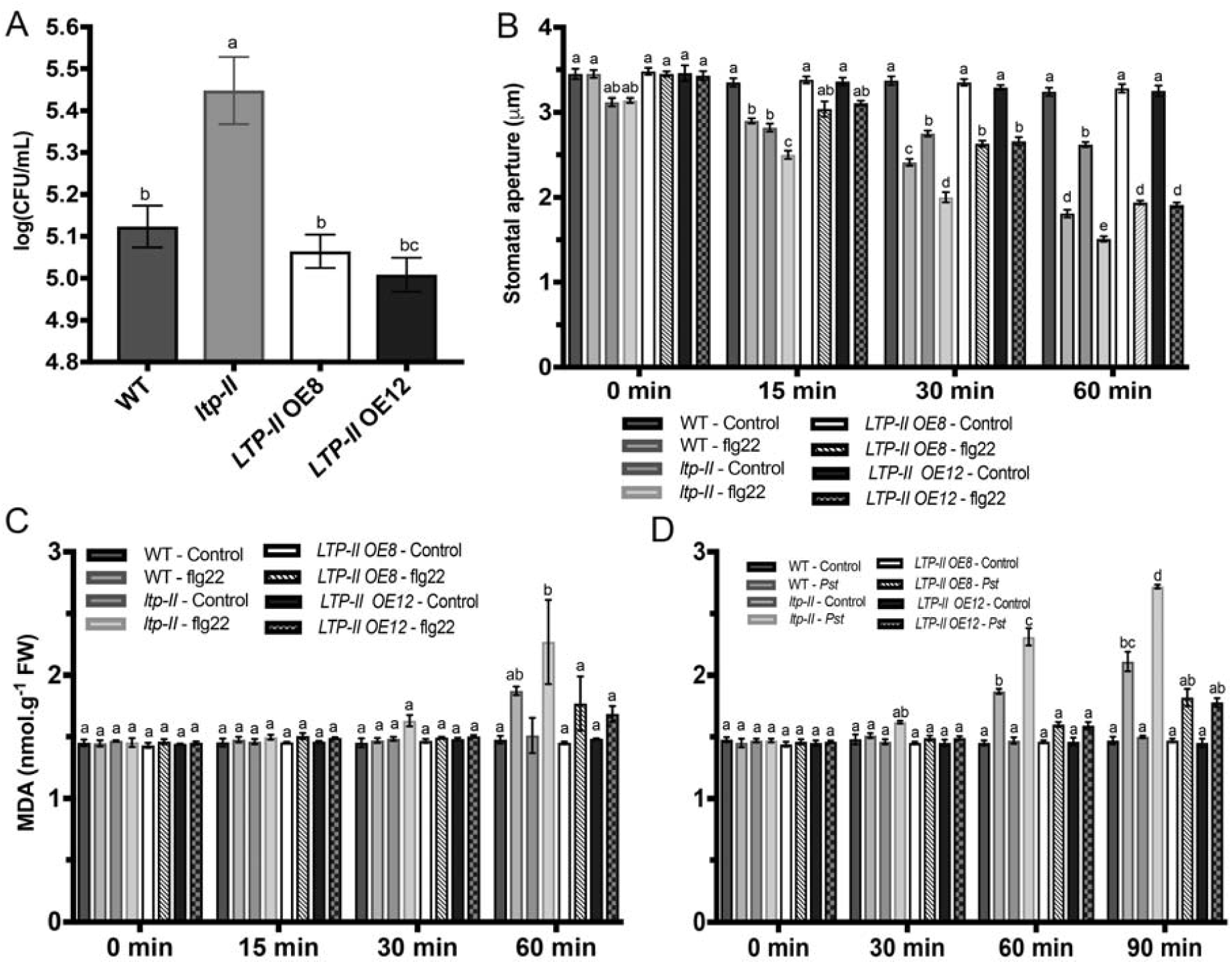
Genetic rescue of the *ltp-II* mutant with *B. napus LTP-II* gene and lipid oxidation in response to flg22 and *Pst* DC3000. (**A**) Bacterial growth in WT, *ltp-II* mutant and two independent overexpressing lines *LTP-OE8* and *LTP-OE12.* The leaves were dip-inoculated with *Pst* DC3000 (OD600= 0.03). Three leaves from each genotype were taken at 3 dpi to determine the bacterial growth. The data are shown as means ± SE of three independent experiments. (**B**) Stomata movement in response to 10μM flg22. Stomatal aperture data were obtained from 180 stomata from three independent experiments and presented as means ± SE. Different letters indicate significantly different mean values at p<0.05. (**C**) Malondialdehyde (MDA) contents in control and 10*μ*M flg22 treated leaves in a time course of 0 min, 15 min, 30 min and 60 min. (**D**) MDA contents in mock control and *Pst* treated leaves. The values were determined in control and 0.05 OD600 treated samples in a time course of 0 min, 15 min, 30 min, 60 min, and 90 min. Data were obtained from three independent experiments and presented as means ± SE.

### *ltp-II* Plants Have Greater Membrane Damage than WT Under Biotic Stress

Increase in cellular ROS levels may result in oxidative stress and lead to membrane damage (Sharma el al., 2012; You and Chan, 2015). To test our hypothesis of LTP-II acting as a ROS scavenger and cellular antioxidant, we assessed the integrity of the cellular membranes under biotic stress (flg22 and *Pst DC3000* treatment) using malondialdehyde (MDA) content, a common indicator of oxidative stress and membrane damage. No significant difference in the *ltp-II* plants was observed at 60 min following flg22 treatment though a slight increase was noticeable (Fig. 4C). In contrast, a significant increase in MDA was observed in the *ltp-II* plants at 60 min after *Pst* DC3000 treatment compared to WT and the *LTP-II* overexpressing lines (Fig. 4D). The observed difference between the *ltp-II* responses to *Pst* DC3000 and flg22 is likely a result of *Pst* DC3000 ability to trigger a robust and more sustained level of ROS in plants. Taken together, the results suggest that the increased susceptibility of *ltp-II* plants to *Pst* may largely be due to oxidative stress leading to loss of membrane integrity and thus plant fitness.

## DISCUSSION

### Redox Regulation and Photosynthesis in Guard Cell Response to Flg22

ROS play important roles in plant defense response, including redox modification of proteins (Mou et al., 2003; Baxter et al., 2014; Balmant et al., 2015; Balmant et al., 2016). In this study, we identified a Cu/Zn SOD being redox-regulated during stomatal closure in response to flg22 (Table 1). SOD is known to play a central role in defense against oxidative stress in plants (Sharma et al., 2012). It catalyzes the dismutation of superoxide radical (O_2_^-^)into O_2_ and H_2_O_2_, and it is present in most of the subcellular compartments that generate oxygen (Sharma et al., 2012). At high concentrations, H_2_O_2_ is able to oxidize and inactivate Cu/Zn-SOD (Sharma et al., 2012). Thus, the reduced form of SOD is active. Here we showed that the Cu/Zn-SOD was reduced at all the time points after flg22 treatment (Tables 3-5, 3-6 and 3-7), indicating that this protein was active in regulating the concentration of O_2_^-^. Interestingly, an aspartic protease in guard cell 2-like was found to undergo oxidation at the late stage of flg22 triggered stomatal closure (Table 1). Yao et al. (2012) demonstrated a role of aspartic protease in controlling ROS levels. Plants overexpressing aspartic proteases were capable of scavenging excessive ROS to prevent oxidative damage through SOD activation (Yao et al., 2012). Another protein, germin-like (GLP) was also observed to be oxidized at the late stage of flg22 triggered stomatal closure (Table 1). GLPs are known to play crucial roles not only in plant development but also in plant defense responses (Wang et al., 2013). It is speculated that the mechanism by which GLPs function in plant defense is associated with their SOD activities (Wang et al., 2013). GLPs are known to be thioredoxin targets (Buchanan and Balmer, 2005) and were found to undergo oxidation in guard cells in response to MeJA (Zhu et al., 2014).

Proteins involved in photosynthesis, including oxygen-evolving enhancer protein1-1, photosystem I reaction center subunit psi-N, RubisCO large subunit, and photosystem I chlorophyll a/b-binding protein 3-1 were also found to be redox-regulated in guard cells in response to flg22 (Table 1). RubisCO large subunit and oxygen-evolving enhancer protein 1 are known to be thioredoxin targets (Lemaire et al., 2004). In addition, ferredoxin has the conserved cysteine residues necessary for thioredoxin-dependent regulation (Walters and Johnson, 2004). Oxidation of these proteins will decrease their activities, supporting the negative impact of biotic stress on photosynthesis.

### Role of LTP-II in the Regulation of ROS Levels in Guard Cells

LTPs encoded by small multigene families are present in many organisms. After more than 40 years of their discovery in plants, none of the LTPs has been extensively characterized. LTPs are small (6.5-10.5 kDa), basic proteins present in high amounts in higher plants (Liu et al., 2015a). It is estimated that they correspond to as much as 4% of the total soluble proteins. In vitro assays have shown the activity of LTPs in transferring phospholipids between membranes as well as binding to acyl chains (Kader, 1996). They are also known to be involved in many other biological processes. For instance, DeBono et al (2009) showed the role of an LTP in wax and cutin metabolism. A mutant of the LTP showed reduced wax load on the stem surface, highlighting the importance of LTP in cuticular lipid export. In addition, an LTP CaMF2 specifically expressed in flower buds of the *Capsicum annuum* male fertile line plays a vital role in pollen development (Chen et al., 2011). Moreover, an LTP OsC6 in rice was demonstrated to play an important role in regulating post-meiotic anther development (Zhang et al., 2010a). A number of studies have also shown that LTPs are involved in cell wall loosening and extension process (Nieuwland et al., 2005; Yeats and Rose, 2008; Liu et al., 2015). This function of LTP in cell wall loosening might contribute to the small stomatal aperture phenotype observed in the *ltp-II* plants (Figs. 2, 4), since wall loosening is an essential step in guard cell swelling (Wei et al., 2011). However, we believe that ROS production and the scavenging capability of LTPII are the main factors for the differences in stomatal aperture between WT and the *ltp-II* plants. LTPs were reported to enable plants to adapt to abiotic stress conditions, such as drought (Guo et al., 2013), cold (Kielbowicz-Matuk et al., 2008), and salt stress (Jang et al., 2004). Last but not least, LTPs are involved in plant defense against biotic stresses, including attack by bacteria, fungi and viruses (Champigny et al., 2013; Liu et al., 2015). For instance, Champigny et al. (2013) identified a lipid transfer protein (DIR1 - Defective in induced resistance 1) as being part of the systemic acquired system (SAR) acting as a long-distance signal molecule. Interestingly, LTPs have been classified as pathogenesis-related (PR) 14 family protein (Sels et al., 2008).

In our study, we identified the LTP-II (protein ID number: 913408) as being oxidized in guard cells in response to flg22, and we didn’t observe significant protein level changes after the flg22 challenge. Although up-regulation of *LTP*s was observed after 3 h, 6 h and 12 h of exposure to flg22, the expression was not changed after 30 min and 1 h of the treatment (Denoux et al., 2008). This result is consistent with our results (Table 1). Regulation of *LTP* expression was also demonstrated in plant responses to cold, drought and ABA treatments (Guo et al., 2013). The expression of *LTP3* was up-regulated after 3 h of treatment, but was not significantly changed upon 1 h of treatment. Although LTPs may share high sequence similarity (Fig. S2), they may have differential cellular and tissue-specific expression patterns and are not completely redundant. Interestingly, this LTP-II is the only LTP identified in the guard cell proteomics experiments (Table 1). The obvious stomatal phenotype and pathogen susceptibility of the *ltp*-II mutant (Figs. 1 and 2) indicate specific functions of the LTP-II.

The oxidation of LTP-II correlates with the peak of ROS production and stomatal closure. Another LTP in *Arabidopsis* cell cultures was also oxidized in response to biotic stress (Liu et al., 2015). Additionally, several studies have reported the role of other LTP family members in controlling ROS levels in plants (Wang et al, 2014; McLaughlin et al, 2015; Xu et al., 2018). Although ROS play important roles in plant defense and stomatal movement (Baxter et al., 2014), over-accumulation of ROS can cause damage to the cells through lipid peroxidation, protein degradation and membrane destruction (Sharma et al., 2012). Furthermore, over-accumulation of ROS (without LTP-II) can cause extreme oxidative stress that can further lead to cellular damage and decrease in plant fitness. Several studies have shown that occurrence of oxidative and abiotic stresses can dramatically alter the response of plants to biotic stress. For instance, hemibiotrophic pathogens can cause severe disease during drought stress. Duniway (1977) showed that drought stress increased the root rot caused by *Phytophthora cryptogea* in *Carthamus tinctorius*. The evidence suggests that the outcome of abiotic stresses and pathogen interaction may lead to increase in severity of diseases in the host plant, which corroborates with our findings. Clearly, over-accumulation of ROS observed in the *ltp-II* plants (Fig. 3), not the small stomatal aperture explains the susceptibility of the *ltp-II* plants to the *Pst* DC3000 infection (Figs. 2, 4).

Based on our results of stomatal movement in the presence of ROS scavenger catalase (Fig. 3), we hypothesize that LTP-II functions as an important ROS scavenger and cellular antioxidant in guard cells. Considering the LTP-II expression patterns in ePlant (Waese et al., 2017), this ROS scavenging function may be more widespread, not limited to guard cells. Additionally, LTP-II may also play a role in downstream plant defense responses similar to antipathogenic peptides, such as DL1 (Kim et al., 2009). Here we proposed a dual mechanism by which LTP-II may function in guard cell immune response and apoplastic defense (Fig. 5). Upon pathogen perception, ROS production is triggered in guard cells, leading to perturbation of cellular redox homeostasis. In this scenario, LTP-II acts as a potent ROS scavenger and becomes oxidized, decreasing the ROS levels to avoid oxidative damage to the cells. The oxidized form of LTP was reported to be the active form of this protein (Kader, 1996). Thus, it can also be speculated that LTP-II can reduce the damage in the membrane by transporting un-damaged lipids from endoplasmic reticulum in the exchange of oxidized membrane lipids. In the meantime, downstream plant defense responses can be activated. When *LTP-II* is knockout, the ROS quenching activity is compromised, leading to membrane damage. Meanwhile, LTP-II lipid transport and repairing activity, as well as plant defense responses are compromised, leading to aggravated damage and thus susceptibility to pathogen infection (Fig. 5). Although our data provide good evidence for such a mechanistic model, detailed microcopic monitoring of LTP-II subcellular relocation, biochemical characterization of LTP-II lipid transport activities, and the functional implication in plant defense are exciting future research directions.

**Fig. 5.**
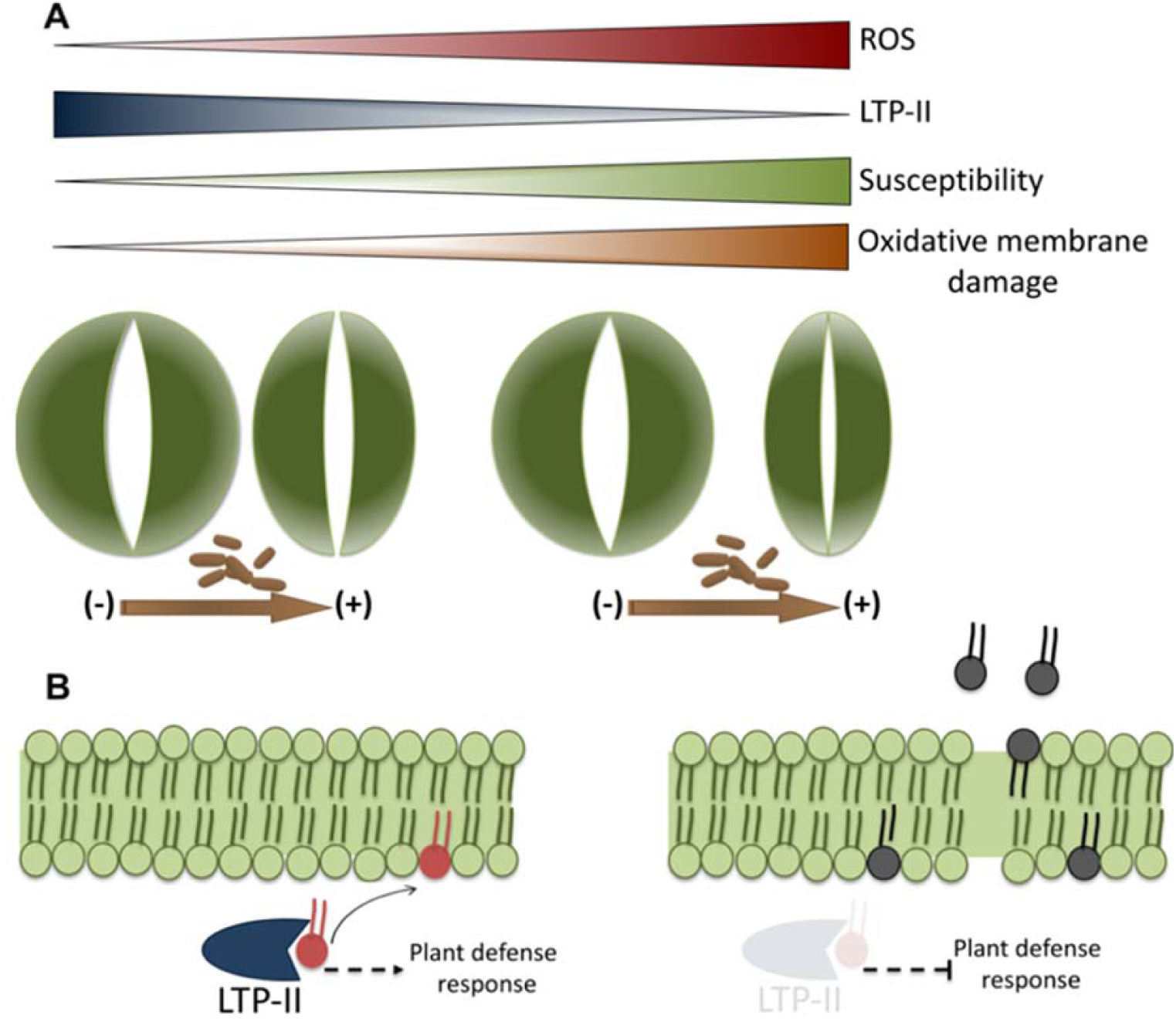
Diagram depicting potential mechanisms underlying the function of LTP-II in stomatal guard cell innate immunity. (**A**) Upon pathogen perception, ROS production is triggered in guard cells, leading to stomatal closure. LTP-II acts as a potent ROS scavenger, reducing the ROS levels to prevent oxidative damage to guard cells. In contrast, the *ltp-II* mutant accumulates more ROS, showing small stomatal aperture and susceptibility to bacterial infection. (**B**) Besides acting as a ROS scavenger, LTP-II also plays a role in stomatal guard cell immunity by repairing the damage in the plasma membrane caused by oxidative stress through its activity in exchanging the oxidized lipids.

## CONCLUSIONS

Uncovering guard cell protein PTM changes in response to stress conditions is necessary for an enhanced understanding guard cell signaling. Using modern proteomic technologies together with genetics, molecular biology, biochemistry and bioinformatics tools, we have shown the utility of MS-based approaches along with biochemical and reverse genetics tools to identify and characterize redox-responsive proteins in the guard cells. This work led to the identification and quantification of dozens of redox-responsive proteins for future hypothesis-testing experiments. More interestingly, it led to the discovery of a lipid transfer protein LTPII regulation via redox and its potential role in plant resistance to bacteria pathogens. The results have expanded our knowledge of guard cell redox signaling and provided strong evidence for the involvement of LTPs in plant defense.

## EXPERIMENTAL PROCEDURES

### Plant Material and Growth Conditions

*Bassica napus* seeds were germinated in moistened soil (Metro-Mix 500 potting mixture from the Scotts Co., USA) and plants were grown in a growth chamber under a photosynthetic flux of 160 µmol photons m^−2^ s^− 1^ with a photoperiod of 8 h light at 22 °C and 16 h dark at 20 °C as previously described (Zhu et al., 2010). *Arabidopsis thaliana* ecotype Columbia (Col-0) and *Arabidopsis LTP-II* T-DNA insertion line (SALK_026257) were obtained from the *Arabidopsis* Biological Resource Center (ABRC, https://abrc.osu.edu). Seeds were germinated on Murashine and Skoog medium in the growth chamber under a photosynthetic flux of 160 µmol photons m^−2^ s^−1^ with a photoperiod of 8 h light at 22 °C and 16 h dark at 20 °C. After 7 days, plant seedlings were transferred to the moistened soil and grown under the same conditions for five weeks. The mutant line was genotyped by amplifying the genomic DNA with left genomic primer (5’-AATATGGTCGATCGTTCCATG-3’) and right genomic primer (5’-GAGGGAAGGAGTAGTTGGTCG-3’). These two genomic primers were used in the same PCR reaction with a T-DNA left border primer (LBa1) as previously described (Yan et al., 2009).

For *Arabidopsis* transformation, the *Arabidopsis ltp-II* mutant was transformed according to an *Agrobacterium*-mediated floral-dip method (Bent, 2008). Briefly *BnLTP-II* was amplified from *B. napus* genomic DNA with primers *BnLTP-II-F* (5’-ATGGCTGGTCTAATGAAGTTG-3’), *BnLTP-II-R* (5’-TTTCACACTGTTGCAGTTGGT-3’) and cloned into pCambia 1305. The construction was transferred to *Agrobacterium* GV3101 which was further used to transform the *ltp-II* mutant plants. Plants overexpression *BnLTP-II* were identified by semi-quantitative PCR with specific primers *LTP-II*-F (5’-CGGTACCCGGGGATCCATGGCTGGTCTAATGAAGTTG -3’) and *LTP-II*-R(5’-AGTCTCCGCCACTAGTTTTCACACTGTTGCAGTTGGT-3’). T4 generation homozygous plants were used for phenotype analysis. All the primers used are listed in Table S2.

### RNA extraction, reverse transcription and PCR

Total RNA was extracted from *A. thaliana* leaves using the RNeasy® Plant Mini Kit (Qiagen, Valencia, CA, USA) following the manufacturer’s protocol. RNA quantity and quality were measured using the NanoDrop® 1000 Spectrometer (Thermo Fisher Scientific, USA) and cDNA was synthesized from 1 µg of total RNA using ProtoScriptR II Reverse Transcriptase (New England BioLabs, MA, USA) in 20 µl reaction with oligo (dT) following the manufacturer’s protocol. *BnLTP-II* primers F- (5’-ATGGCTGGTCTAATGAAGTTG-3’) and R- (5’-TTTCACACTGTTGCAGTTGGT-3’) were used in a PCR reaction with the newly synthesized cDNA. PCR was preformed using Taq DNA Polymerase (New England Biolabs, Beverly, MA, USA) as follows: 98 °C for 30 s (1 cycle); 98 °C for 10 s, 55 °C for 30 s, and 72 °C for 30 s (30 cycles); and 72 °C for 10 min (1 cycle).

### Guard Cell Enrichment, Flg22 Treatment and Stomatal Movement Assay

Fully expanded leaves from 7 week old *B. napus* were used for the preparation of enriched guard cells as previous described (Zhu et al., 2016). After guard cell enrichment, the samples were left for recovery for 1 h under the same conditions in the growth chambers described above. Enriched stomatal samples were assessed for guard cell purity and viability using neutral red and fluorescein diacetate (FDA), respectively. For flg22 treatment, guard cell enriched samples were incubated with 10 μM flg22 under gentle shaking (30 rpm) for 15 min, 30 min and 60 min.

Stomatal aperture measurement was carried out as previously described (Zhu et al., 2010) with slight modifications. Leaves from 7-week-old *B. napus* and 5-week-old *A. thaliana* were blended, and the epidermal peels were washed with water. The epidermal peels were subjected to enzymatic digestion to obtain enriched stomatal samples. The freshly prepared epidermal peels and enriched stomatal samples were incubated in a degassed opening medium (50 μM CaCl_2_, 10 mM KCl and 10 mM MES-KOH, pH 6.2) for 2 h under light to promote stomata opening. In the case of *Arabidopsis* epidermal peels, the degassed medium was supplemented with 20 μM propidium iodide (PI) to stain the cell wall of guard cells and facilitate the stomatal aperture measurement. After checking the stomatal opening, 1 μM, 3 μM and 10 μM flg22 or 200 U/mL catalase were added into the samples. For both experiments, at the indicated time points, images of stomata were captured at a 200X magnification using a Zeiss Axiostar Plus microscope (Carl Zeiss Inc., USA). Stomatal apertures of at least 50 stomata were analyzed in each experiment and three biological replicates were performed. Stomatal apertures were measured by Image J (NIH, MD, USA) analysis of the digital images. All results were presented as means ± standard error of the three replicates. Data were analyzed using one-way ANOVA followed by Tukey’s test.

### ROS Production and Free Thiol Assays

ROS production in *B. napus* stomatal guard cells was quantified using an OxiSelect Hydrogen Peroxide Assay Kit (Fluorometric - CELL BioLABS, Inc., San Diego, CA), according to manufacturer’s instruction. Briefly, enriched stomatal samples were incubated in the degassed opening medium for 1 h under light to promote stomatal opening and to allow cells to recover from potential stress caused during epidermal cells digestion. After this incubation step, 10 μM flg22 was added to three independent samples. Water was used as mock control. At different time points, an aliquot of 500 μL was removed and centrifuged at 10,000 rpm for 5 min to remove debris. Aliquots of 50 μL supernatant were mixed with 50 μL of ADHP/HRP (10-Acetyl-3, 7-dihydroxyphenoxazine/ horseradish peroxidase) working solution and added into fluorescence compatible microtiter plate. After incubation at room temperature for 30 min in the dark, the plate was analyzed at 595 nm to obtain the relative fluorescence unit values. Results were presented as means ± standard error of three replicates. Data were analyzed using one-way ANOVA followed by Tukey’s test.

ROS production in *A. thaliana* was examined using the redox-sensitive 2’-7’-dihydro-dichlorofluorescein diacetate (H_2_DCF-DA). Briefly, a couple of leaves were blended and the epidermal peels were incubated in the degassed opening medium for 1 h under light. The epidermal peels were then placed into a buffer containing 50 mM Tris-KCl (pH 7.2) supplemented with 50 μM of H_2_DCF-DA, and incubated for 30 min. Images of stomata were captured using the Zeiss Axiostar Plus microscope with excitation 450-490 nm and emission 520-560 nm. The fluorescence emission levels of at least 150 stomata from three independent experiments were analyzed using Image J (NIH, MD, USA).

Free thiols from stomatal samples were measured using 5, 5’-dithio-*bis*-(2-nitrobenzoic acid) (DTNB) according to manufacture instructions. Epidermal peels of control and flg22-treated plants were ground in liquid nitrogen. Samples were suspended in 1.5 mL of extraction buffer (50 mM Tris-HCl, pH 7.5, 0.1 mM EDTA, 0.1% SDS), and centrifuged at 10,000 g for 15 min at 4°C. A total of 20 μg of protein for each sample was used to measured free thiols.

### Protein Extraction, CysTMTRAQ, Strong Cation Exchange and Nanoflow LC-MS/MS

Three enriched stomatal preparations were pooled to yield one biological replicate. The enriched stomatal samples were ground in liquid nitrogen and protein extraction was carried out as previously described (Parker *et al.*, 2015), and 100 μg of protein (measured by a EZQ protein quantification kit) was used for the cysTMTRAQ double labeling as described by Parker *et al.* (2015). Three control biological replicates were labeled with cysTMT tags (126 *m*/*z*, 128 *m*/*z*, and 130 *m*/*z*) and then with iTRAQ tags (114 *m*/*z*, 117 *m*/*z*, and 119 *m*/*z*). The three flg22-treated biological replicates were labeled with cysTMT tags (127 *m*/*z*, 129 *m*/*z*, and 131 *m*/*z*) and then with iTRAQ tags (115 *m*/*z*, 116 *m*/*z*, and 121 *m*/*z*).

Peptides were dissolved in strong cation exchange (SCX) solvent A (25% (v/v) acetonitrile, 10 mM ammonium formate, and 0.1% (v/v) formic acid, pH 2.8) and fractionated using an Agilent HPLC 1260 system with a polysulfoethyl A column (2.1 × 100 mm, 5 μm, 300 Å). Peptides were eluted at a flow rate of 300 µL/min with a linear gradient of 0–20% solvent B (25% (v/v) acetonitrile and 500 mM ammonium formate, pH6.8) over 80 min, followed by ramping up to 95% solvent B in 5min. Peptide absorbance at 280 nm was monitored, and a total of 12 fractions were collected. For a 125 min nanoflow LC separation of the SCX fractions on a Thermo Scientific EASY-nLC 1000, the flow was 300 nL/min and the column was a 25 cm EasySpray PepMap100 (Thermo Scientific Inc., Bremen, Germany). The gradient conditions were as follow: 5–35% B over 120 min, 35–95% over 1 min, and then held at 95% B for 4 min. MS/MS analysis was carried out on an Orbitrap Fusion mass spectrometer in a positive mode applying data-dependent MS and MS/MS acquisition (Thermo Scientific, Bremen, Germany). A full scan was performed at 120,000 resolution, scanning from 400 m/z to 1800 m/z with a target of 200,000 ions in the Orbitrap a 50 ms maximum injection time. MS/MS scanning was performed at 60,000 resolution with a quadrupole isolation width of 2 m/z, using a fixed first mass of 100 m/z. The MS/MS target was 50,000 ions with a 70 ms maximum injection time. Ions with charge states 2 to 7 were sequentially fragmented by high-energy collisional dissociation (HCD) with a normalized collision energy (NCE) of 40%. The dynamic exclusion duration was set as 30 s.

### Database Searching and Data Analysis

Proteome Discoverer 1.4 with the SEQUEST algorithm (Thermo Scientific Inc., Bremen, Germany) was used to search a green plants database from NCBI (5,222,402 entries). Proteome Discoverer nodes for spectrum grouper and spectrum selector were set to default with the spectrum properties filter set to a minimum and maximum precursor mass of 300 Da and 5 kDa, respectively. The SEQUEST algorithm was used for protein identification. Parameters were set to two maximum missed cleavage sites of trypsin digestion, absolute XCorr threshold of 0.4, and fragment ion cutoff percentage at 0.1. Tolerances were set to a 10-ppm precursor mass tolerance and a 0.01 Da fragment mass tolerance. Percolator was used for protein identification with parameters of a strict target false discovery rate of 0.01 and a relaxed target false discovery rate of 0.05. Quantification was performed using the reporter ion peak areas of both cysTMT and iTRAQ. Peptide results were filtered to include only peptides with high confidence of identification (1% false discovery rate, FDR).

After peptide and protein identification, fold changes were compared among the respective reporter ions of iTRAQ and cysTMT. The median values of the peak areas were used to normalize the data. The reporter ion peak areas of identical peptides were summed, log2 transformed, and the ratios between control and treated samples were obtained. For protein quantification, at least three unique peptides were required. Protein quantification was performed by summing the reporter ion peak areas of all corresponding peptides (Carrillo et al., 2010), and ratios were generated as described above. Student’s *t* test (two-tail) on the log2-transformed treated/control ratios was performed, and the *p* values were further corrected via the q-value FDR method (Storey, 2002). A peptide with a q-value less than 0.05 and a fold change greater than 1.2 or less than 0.8 was considered statistically significant. The significant peptides labeled with cysTMT were compared with the significant proteins quantified via iTRAQ. Student’s *t* test was conducted between the fold changes of cysTMT-labeled peptides and the fold changes of the corresponding proteins quantified via iTRAQ. A correction factor was applied to the fold change of cysTMT-labeled peptides, taking into account the fold change of the protein quantified via iTRAQ.

### Bacterial Growth Assay

The bacteria growth assay was carried out as previously described (Yao et al., 2013). Briefly, a single colony of *Pseudomonas syringae* pv. *tomato* DC3000 (*Pst* DC3000) was grown on Kings B Medium (KBM) at 28°C overnight. The bacteria inoculum was prepared by scraping the bacterial colony from the KBM plate into a solution containing 0.02% Silwet L-77 at a concentration of 1×10^8^ CFU/mL. The mock control samples were inoculated with the same solution but without bacteria. Five weeks old *A. thaliana* plants (WT, *ltp-II* mutant, *LTP-OE8* and *LTP-OE12*) were inoculated by dipping into the above inoculation solution for 5 seconds. The plants were covered and placed in the growth chamber for 3 days. A disc (1cm^2^) was obtained from each leaf sample and ground in 10 mM MgCl_2_. The solution was diluted and spotted onto a KBM Rifampicin (25 mg/ml) and Kanamycin (50 mg/mL) plate. Colonies were counted, and a t-test was performed to determine whether there are differences in bacterial growth among WT, the *ltp-II* mutant and the overexpressing lines. The experiments were repeated three times.

### Determination of MDA Content

Leaves of 5-week-old *Arabidopsis* plants were weighed and homogenized in 1mL of 10% trichloroacetic acid (TCA) using a mortar and pestle. The samples were centrifuged for 10 min at 10,000rpm and the supernatant was added to a microcentrifuge tube containing a mixture of 0.6% thiobarbituric acid in 10% TCA. The mixture was placed in boiling water and incubated for 30 min. The reaction was stopped by placing the mixture on ice for 3 min. The samples were centrifuged, and the absorbance of the supernatant was measured at 450 nm, 532 nm and 600 nm. MDA contents (nmol g^-1^ fresh weight) were calculated using the formula [6.45(A_532_ − A_600_) − 0.56A_450_]/ fresh weight.

## ACKNOWLEDGEMENTS

This work was supported by the U.S. National Science Foundation grants NSF-0818051 and NSF- 1412547 to SC.

